# RankMap: Rank-based reference mapping for fast and robust cell type annotation in spatial and single-cell transcriptomics

**DOI:** 10.64898/2026.03.01.708931

**Authors:** Jinming Cheng, Shengdi Li, Serim Kim, Chow Hiang Ang, Sin Chi Chew, Pierce Kah Hoe Chow, Nan Liu

## Abstract

Accurate cell type annotation is essential for the analysis of single-cell and spatial transcriptomics data. While reference-based annotation methods have been widely adopted, many existing approaches rely on full-transcriptome profiles and incur substantial computational cost, limiting their applicability to large-scale spatial datasets and platforms with partial gene panels. Here, we present *RankMap* (https://github.com/jinming-cheng/RankMap), an efficient and flexible R package for reference-based cell type annotation across both single-cell and spatial transcriptomics. *RankMap* transforms gene expression profiles into rank-based representations using the top expressed genes per cell, improving robustness to platform-specific biases and expression scale differences. A multinomial regression model trained with elastic net regularization is then used to predict cell types and associated confidence scores. We benchmarked *RankMap* on five spatial transcriptomics datasets, including Xenium, MERFISH, and Stereo-seq, as well as two single-cell datasets, and compared it with established methods such as *SingleR, Azimuth*, and *RCTD. RankMap* achieved competitive or superior annotation accuracy while consistently reducing runtime compared to existing methods, particularly for large spatial datasets. These results demonstrate that *RankMap* provides a scalable and robust solution for reference-based cell type annotation in modern single-cell and spatial transcriptomics studies.

## 1 Introduction

Accurate cell type annotation is a foundational step in the analysis of single-cell and spatial transcriptomics data. In single-cell RNA sequencing (scRNA-seq) and single-nucleus RNA sequencing (snRNA-seq), reference-based annotation methods are widely used to map query cells to well-annotated reference atlases, enabling systematic comparisons across datasets and studies.

With the growing availability of high-resolution spatial transcriptomics platforms, including Xenium [1], MERFISH [2], Stereo-seq [3], CosMx [4] and STARmap [5], that provide spatially resolved gene expression at or near single-cell resolution, there is a pressing need for annotation tools that are accurate, scalable, and robust to platform-specific technical variability.

Numerous methods have been developed for cell type deconvolution or label transfer in spatial transcriptomics, such as *Seurat* ‘s label transfer [6], *RCTD* [7], *Cell2location* [8], and *Tangram* [9]. While effective in many contexts, these methods often require whole-transcriptome coverage, complex hyperparameter tuning, or substantial computational resources. This can limit their applicability to emerging platforms with partial gene panels, such as Xenium and MERFISH, or large datasets containing hundreds of thousands of spatially resolved cells.

Several reference-based annotation methods, including *SingleR* [10] and *Azimuth* [11] for single-cell data, as well as *RCTD* [7] for spatial transcriptomics, have demonstrated strong performance on Xenium datasets [12]. However, these methods can face challenges when applied to large-scale or heterogeneous spatial datasets. These include high memory and runtime requirements, reduced robustness to sparsity and noise, and difficulties generalizing across spatial technologies with differing expression quantification protocols.

To address these challenges, we developed *RankMap*, a fast and flexible R package for reference-based cell type annotation based on rank-transformed gene expression. Rather than relying on raw or normalized expression values, *RankMap* leverages the ranks of the top `k` expressed genes per cell, enhancing robustness to batch effects and cross-platform differences. The ranked matrix can be optionally refined via binning, expression weighting, and normalization to emphasize informative features and reduce variance artifacts. Cell type prediction is performed using multinomial logistic regression implemented via the *glmnet* [13] framework, enabling efficient training and scalable inference.

*RankMap* supports both single-cell and spatial transcriptomics input formats and is compatible with common R data structures such as `Seurat`, `SingleCellExperiment`, and `SpatialExperiment`. Its streamlined design enables fast training and prediction, making it well suited for large-scale datasets, though memory usage may still be a consideration for very large inputs.

In this study, we benchmark *RankMap* against existing reference-based methods across five spatial transcriptomics datasets and two single-cell transcriptomics collections. These datasets span multiple technologies (Xenium, MERFISH, Stereo-seq, scRNA-seq, and snRNA-seq) and diverse tissues (mouse brain, human breast cancer, human lung, macaque cortex, and human liver), providing a comprehensive evaluation across biological and technical contexts. Our results demonstrate that *RankMap* achieves competitive or superior annotation accuracy while offering substantial improvements in runtime and scalability.

Together, these findings position *RankMap* as a robust, efficient, and user-friendly framework for reference-based cell type annotation across spatial and single-cell transcriptomics, enabling reproducible and high-throughput analysis in the era of large-scale spatial biology.

## 2. Methods

### 2.1 Mathematical Formulation of RankMap

#### 2.1.1 Rank Transformation of the Expression Matrix

Let *X* ∈ ℝ^*G×N*^ denote the log-normalized gene expression matrix, where *G* is the number of genes and *N* is the number of cells or spatial spots. For each column *X*_·,*n*_, *RankMap* retains only the top *k* expressed genes and assigns them a rank based on expression magnitude:

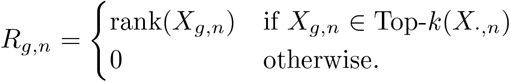

After this sparsified ranking, *R* ∈ ℝ^*G×N*^ can undergo additional transformations to improve its utility for classification. First, rank values may be discretized into a fixed number of equal-width bins, reducing sensitivity to small expression differences. Second, each (binned) rank may be multiplied by log(1 + *X*_*g,n*_) to incorporate expression magnitude. Finally, gene-wise z-score standardization can be applied to normalize variance across cells. These three transformations—binning, weighting by expression magnitude, and scaling—are enabled by default, as this configuration produced the best overall performance in our spatial transcriptomics benchmarking (Xenium datasets).

We denote the resulting transformed matrix as 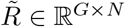.

#### 2.1.2 Multinomial Classification Model

Let *y*_*n*_ ∈ {1, …, *C*} denote the cell type label for sample *n. RankMap* employs a multinomial logistic regression model to estimate class probabilities from the transformed feature vector 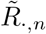:

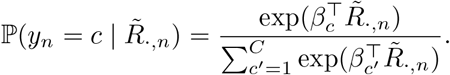

Here, *β*_*c*_ ∈ ℝ^*G*^ is the coefficient vector associated with cell type *c*. The coefficients are estimated by minimizing a penalized negative log-likelihood with elastic net regularization:

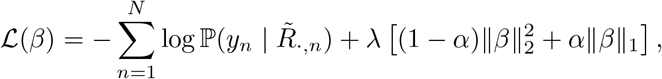

where *λ* controls the overall degree of regularization and *α* ∈ [0, 1] governs the balance between ridge *(ℓ*_2_) and lasso (*ℓ*_1_) penalties. Optimization is performed using the `glmnet` package.

#### 2.1.3 Prediction and Confidence Scoring

For a new sample with features 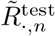, the predicted label is obtained as:

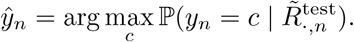

*RankMap* additionally reports a confidence score, defined as the maximum predicted probability:

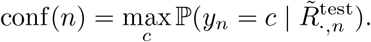

A user-defined threshold *τ* may be applied to filter low-confidence predictions:

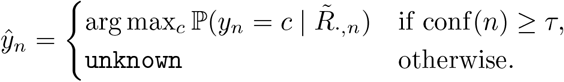

This filtering step enables users to identify ambiguous cells or spots and restrict analyses to high-confidence annotations.

### 2.2 Spatial and single-cell transcriptomics datasets

We used a diverse set of spatial and single-cell transcriptomics datasets for benchmarking *RankMap* across multiple tissue types, technologies, and species:

1. **Mouse brain (Xenium spatial)**. A processed Xenium dataset of fresh-frozen mouse brain tissue was obtained from Cheng et al. [14]. The raw data is publicly available from 10x Genomics at: https://www.10xgenomics.com/datasetsfresh-frozen-mouse-brain-for-xenium-explorer-demo-1-standard.
2. **Mouse brain (scRNA-seq)**. A 10,000-cell subset of the mouse brain single-cell dataset previously reanalyzed by Cheng et al. [14] was used as the reference for the mouse brain Xenium data. The full processed dataset originates from the Allen Brain study [15] and can be accessed via GEO under accession: GSE185862.
3. **Human HER2+ breast cancer (Xenium and snRNA-seq)**. A matched processed Xenium and snRNA-seq dataset from a human HER2+ breast tumor (Sample 1) was obtained from Cheng et al. [12], along with their curated cell type annotations: https://github.com/jinming-cheng/cheng_annotation_bmc_bioinfo. A 5,000-cell subset of the snRNA-seq data was used as reference. Raw data is available from 10x Genomics: https://www.10xgenomics.com/products/xenium-in-situpreview-dataset-human-breast[1].
4. **Human ER+ breast cancer (scRNA-seq)**. Twelve ER+ breast cancer scRNA-seq samples were downloaded from GEO under accession: GSE161529 [16]. These samples were reanalyzed in this study, using sample ER_Total_0360 as the reference for cell type annotation.
5. **Human lung (Xenium spatial)**. The human lung Xenium dataset was downloaded from 10x Genomics: https://www.10xgenomics.com/datasetsxenium-human-lung-preview-data-1-standard.
6. **Human lung (scRNA-seq)**. Healthy lung scRNA-seq samples were obtained from the Chan Zuckerberg CellxGene portal [17]: https://cellxgene.cziscience.comcollections/6f6d381a-7701-4781-935c-db10d30de293. Eight samples (Donor 01–08) were selected for analysis, with Donor 02 used as the reference for both the Xenium and scRNA-seq benchmarking.
7. **Macaque cortex (Stereo-seq and snRNA-seq)**. The raw count matrix and annotations for a Stereo-seq dataset (slide T33) were obtained from the authors [18], while the matched snRNA-seq dataset was downloaded from: https://www.braindatacenter.cn/datacenter/web/#/dataSet/details?id=1663381185152036865. Two samples (101T99-1-210421 and 101T99-2-210421) were extracted and used as the single-cell reference.
8. **Human liver (MERFISH and snRNA-seq)**. A MERFISH dataset of healthy human liver tissue, along with a matched snRNA-seq dataset, was downloaded from Dryad [19]: https://datadryad.org/stash/dataset/doi:10.5061/dryad.37pvmcvsg. Subsets of sample AM042 were used for both spatial and reference modalities in our analysis.

### 2.3 Manual annotation of single-cell and spatial transcriptomics datasets

Manually curated or author-provided annotations were used as the ground truth for benchmarking cell type prediction performance. For each spatial dataset, corresponding single-cell or single-nucleus reference data were first annotated or refined before transferring labels to the spatial data.

For the mouse brain scRNA-seq dataset, author-provided cell type labels were manually merged to match the simplified taxonomy used by Cheng et al. [14]. For the HER2+ human breast cancer snRNA-seq data, we used the cell type annotations from Cheng et al. [12] without modification. For the human lung scRNA-seq dataset, author-provided annotations were used directly.

The macaque cortex snRNA-seq data were manually re-annotated by merging fine-grained labels into broader categories. Specifically, layer-specific excitatory neuron types were grouped into “L2/3/4” (including L2, L2/3, L2/3/4, L3/4, L3/4/5) and “L4/5/6” (including L4, L4/5, L4/5/6, L5/6). Similarly, interneuron subtypes VIP and VIP_RELN were combined into a single “VIP” category. For the human liver MERFISH dataset, macrophage subtypes (Mac_1 and Mac_2) were merged into “Mac”, and hepatic stellate cell subtypes (HSC_1 and HSC_2) into “HSC”.

The human ER+ breast cancer scRNA-seq dataset was reprocessed using the *Seurat* (v5.3.1 and v5.1.0) pipeline [6]. Standard preprocessing steps included quality control (filtering low-quality cells), normalization (`NormalizeData`), identification of variable genes (`FindVariableFeatures`), data scaling (`ScaleData`), dimensionality reduction (`RunPCA`, `RunUMAP`), and graph-based clustering (`FindNeighbors`, `FindClusters`). Resulting clusters were manually annotated based on canonical marker gene expression.

The *Seurat* pipeline was also applied to all spatial transcriptomics datasets for consistency. For the mouse brain and human lung Xenium datasets, cluster-level annotations were guided by marker genes identified from the corresponding single-cell reference using `FindAllMarkers`. Clusters were assigned to the most likely cell type based on their marker expression profiles. The annotations from Cheng et al. [12] were used directly for the HER2+ breast cancer Xenium data. For the macaque cortex Stereo-seq and human liver MERFISH datasets, the spatial annotations were manually merged in the same way as the single-cell references described above.

### 2.4 Benchmarking against existing annotation methods

For spatial transcriptomics datasets, we benchmarked *RankMap* against *glmnet_expr* (implemented within the *RankMap* framework) and three commonly used reference-based annotation methods: *SingleR* (v2.8.0) [10], *Azimuth* (v0.5.0) [11], and *RCTD* (*spacexr* v1.0.0) [7]. Each spatial dataset was paired with a corresponding single-cell reference dataset, as described above.

For single-cell transcriptomics datasets, we compared *RankMap* to *SingleR* and *Azimuth* across twelve human ER+ breast cancer samples and eight healthy human lung samples. ER_Total_0360 and Donor_02 were used as reference samples for the breast and lung datasets, respectively.

Benchmarking was performed using the set of genes shared between the query dataset and its reference. Prediction accuracy was computed as the proportion of query cells with predicted labels matching the ground truth. Runtime was measured using `Sys.time()` in R, focusing on the core annotation functions: `RankMap()`, `SingleR()`, `RunAzimuth()`, and `runRctd()`.

### 2.5 Preprocessing and method-specific settings

For *RankMap* and *glmnet expr*, both query and reference datasets were provided as `Seurat` objects. In *RankMap*, the `k` parameter was set to 100, 20, 100, 100, and 20 for the mouse brain Xenium, human HER2+ breast cancer Xenium, human lung Xenium, macaque cortex Stereo-seq, and human liver MERFISH datasets, respectively. For single-cell benchmarking, `k = 300` was used. In the *glmnet_expr* variant, the ranking step was bypassed by setting `use_data = TRUE`, allowing the model to use normalized expression directly.

For *SingleR*, both query and reference datasets were formatted as `SingleCellExperiment` objects.

*Azimuth* required a reference preparation step using the `AzimuthReference` function. Prior to this, we ran `SCTransform` and `RunUMAP(return.model = TRUE)` on the reference data. Due to memory requirements, the system stack size was increased via `ulimit-s unlimited`. Query datasets were passed as `Seurat` objects, and reference files were loaded from a precomputed directory when calling `RunAzimuth()`.

For *RCTD*, a `SpatialExperiment` object (query spatial data) and `SingleCellExperiment` object (reference data) were used as inputs to `createRctd()`. To avoid filtering low-UMI spots or genes, parameters were adjusted as follows: `UMI_min = 0`, `gene_cutoff = 0`, `fc_cutoff = 0`, `fc_cutoff_reg = 0`, `UMI_min_sigma = 1`, `pixel_count_min = 1`, `ref_UMI_min = 10`, and `ref_n_cells_min = 2`. Cell type deconvolution was then performed using `runRctd()` with the default doublet mode. The top predicted cell type per spot was used as the final output.

By default, *RankMap, SingleR*, and *Azimuth* were run using a single CPU, while *RCTD* was executed using four CPUs. Spatial benchmarking was conducted on an AWS EC2 Ubuntu virtual machine with 128 GB of RAM. Single-cell benchmarking was performed on a MacBook Pro with 16 GB of RAM.

### 2.6 Tuning of the k parameter in RankMap

The `k` parameter in *RankMap* determines the number of top-ranked genes retained per cell and is a key hyperparameter influencing performance. In practice, optimal `k` values may depend on dataset characteristics, such as transcriptome coverage or the number of distinct cell types.

To assess the impact of `k`, we systematically evaluated *RankMap* performance using a range of values. For the macaque cortex Stereo-seq dataset (whole-transcriptome), we tested `k = 100, 200, 300, 400, 500, 600`. For the other four spatial datasets (with targeted gene panels), we tested `k = 20, 30, 40, 50, 100, 150, 200`. Accuracy was computed for each setting to assess sensitivity to `k` and identify optimal values for downstream analyses.

## 3 Results

### 3.1 Overview of the RankMap pipeline and benchmarking strategy

To enable scalable and accurate cell type annotation across diverse single-cell and spatial transcriptomics platforms, we developed *RankMap*, a rank-based reference mapping pipeline (Figure 1a). *RankMap* begins with a well-annotated single-cell reference dataset. For each cell, the top-`k` highly expressed genes are selected and ranked, producing a ranked matrix that encodes expression order rather than magnitude. This transformation improves robustness to platform-specific biases and sparsity, particularly in spatial data.

**Fig. 1.**
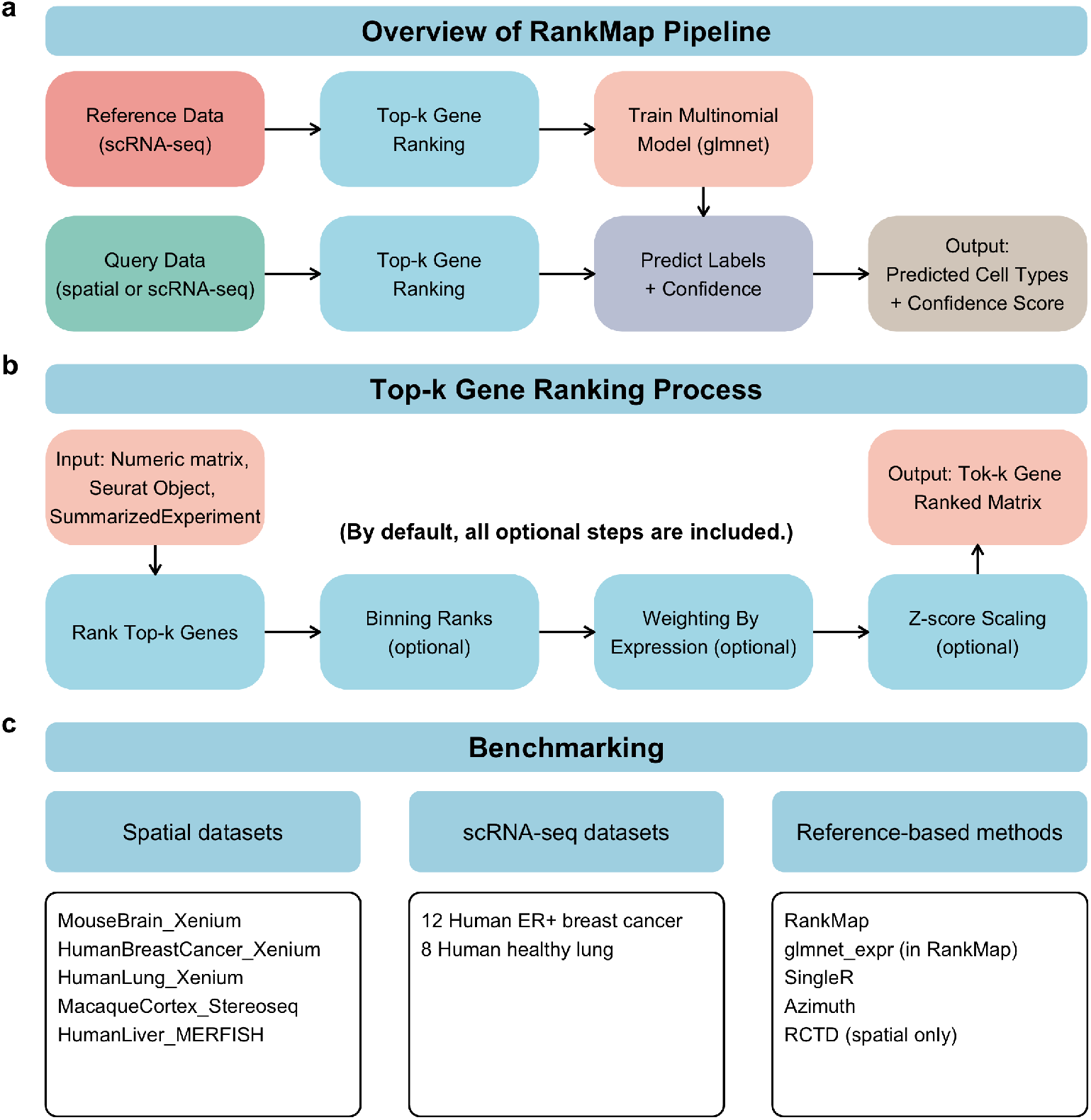
Overview of the RankMap pipeline and benchmarking analysis. **a.**Schematic of the RankMap pipeline. A well-annotated scRNA-seq dataset is used as a reference. For each cell, the top-*k* expressed genes are retained and ranked to construct a ranked matrix. This matrix is transformed and used to train a multinomial regression model using `glmnet`. The trained model is then applied to new scRNA-seq or spatial transcriptomics data using the same top-*k* ranking procedure, generating predicted cell types and confidence scores. **b**. Illustration of the top-*k* gene ranking and transformation process. The input can be a numeric matrix, a Seurat object, or a `SummarizedExperiment`-derived object (e.g., `SingleCellExperiment`, `SpatialExperiment`). The top-*k* genes per cell are retained while others are masked. Ranks are optionally binned into equal-width intervals, then optionally weighted by log(1 + expression) to emphasize high-expression genes. Finally, the matrix is optionally z-score scaled across cells to normalize variance. By default, all three steps (binning, weighting, scaling) are applied before modeling. **c**. Summary of datasets and reference-based methods used in benchmarking. For spatial transcriptomics, we benchmarked three Xenium datasets (mouse brain, human breast cancer, human lung), one Stereo-seq dataset (macaque cortex), and one MERFISH dataset (human liver). For scRNA-seq, we used twelve ER+ breast cancer samples and eight healthy lung samples. RankMap was benchmarked against `glmnet` (direct use of normalized expression), SingleR, Azimuth, and RCTD (spatial only). For single-cell datasets, Azimuth and SingleR were used as benchmarks.

To further enhance signal and standardize inputs across datasets, *RankMap* optionally performs three transformation steps on the ranked matrix: (1) binning ranks into equal-width intervals, (2) weighting ranks by log-transformed expression, and (3) z-score normalization across cells to adjust for variance scale (Figure 1b). These steps are applied by default and can be customized depending on the data modality and use case. The transformed matrix is then used to train a multinomial logistic regression model using the *glmnet* framework, resulting in a lightweight and interpretable classifier.

Once trained, the model can be applied to new spatial or single-cell datasets using the same top-`k` ranking procedure to predict cell type labels and confidence scores. *RankMap* is compatible with common R data structures, including `Seurat`, `SingleCellExperiment`, and `SpatialExperiment`, allowing seamless integration into existing single-cell and spatial analysis workflows.

To assess the performance of *RankMap*, we conducted benchmarking analyses on five spatial datasets and two single-cell datasets (Figure 1c, Supplementary Table 1). The spatial datasets span a range of technologies and species, including Xenium (mouse brain, human breast cancer, human lung), Stereo-seq (macaque cortex), and MERFISH (human liver). The single-cell benchmarking included twelve ER+ breast cancer samples and eight healthy lung samples. *RankMap* was compared against *glm-net expr* (which uses normalized expression instead of ranks), as well as commonly used reference-based annotation methods: *SingleR, Azimuth*, and *RCTD* (spatial only). For single-cell datasets, we benchmarked against *Azimuth* and *SingleR*.

Together, these benchmarking analyses demonstrate the broad applicability of *RankMap* across tissues, species, and data modalities, while highlighting its computational efficiency and competitive accuracy relative to existing tools.

### 3.2 Benchmarking RankMap on spatial transcriptomics datasets

To evaluate the performance of *RankMap* on spatial transcriptomics data, we bench-marked it against four methods: *glmnet_expr* (a variant of *RankMap* that uses normalized expression values directly), *SingleR, Azimuth*, and *RCTD*. We performed this benchmarking across five spatial datasets representing diverse tissues and platforms: three Xenium datasets (mouse brain, human HER2+ breast cancer, human lung), one Stereo-seq dataset (macaque cortex), and one MERFISH dataset (human liver).

Ground-truth cell type annotations were derived from manual annotations or author-provided labels. For the Xenium datasets, annotations were curated based on reference single-cell data and marker gene profiles [12, 14]. The macaque cortex Stereo-seq dataset used author annotations generated via *Spatial-ID* [18, 20], while the MERFISH liver dataset incorporated annotations based on both marker genes and label-transfer [19].

To ensure fair comparison, we used only the set of genes shared between each spatial dataset and its corresponding reference single-cell dataset. The number of common genes and query cells/spots were as follows: mouse brain Xenium (247 genes, 36,402 cells), human breast cancer Xenium (310 genes, 159,226 cells), human lung Xenium (392 genes, 288,837 cells), macaque cortex Stereo-seq (15,759 genes, 393,837 cells), and human liver MERFISH (369 genes, 44,963 cells). The corresponding reference datasets contained 10,000 cells (mouse brain), 5,000 cells (breast cancer), 3,832 cells (lung), 6,892 cells (macaque cortex), and 3,851 cells (liver).

Prediction accuracy was defined as the proportion of query cells/spots whose predicted label matched the ground-truth annotation. Across the five spatial datasets, *RankMap* achieved accuracy values of 0.70 (mouse brain), 0.66 (breast cancer), 0.52 (lung), 0.50 (macaque cortex), and 0.53 (liver), with an average accuracy of 0.582 (Figure 2a). These values were comparable to those of *Azimuth* (0.77, 0.71, 0.55, 0.41, 0.49; average 0.586) and *RCTD* (0.71, 0.72, 0.51, 0.49, 0.48; average 0.582), and higher than *SingleR* (0.67, 0.70, 0.60, 0.50, 0.33; average 0.560) and *glmnet_expr* (0.74, 0.59, 0.56, 0.36, 0.39; average 0.528). Notably, while *glmnet_expr* sometimes achieved relatively high accuracy, its performance was inconsistent across datasets.

**Fig. 2.**
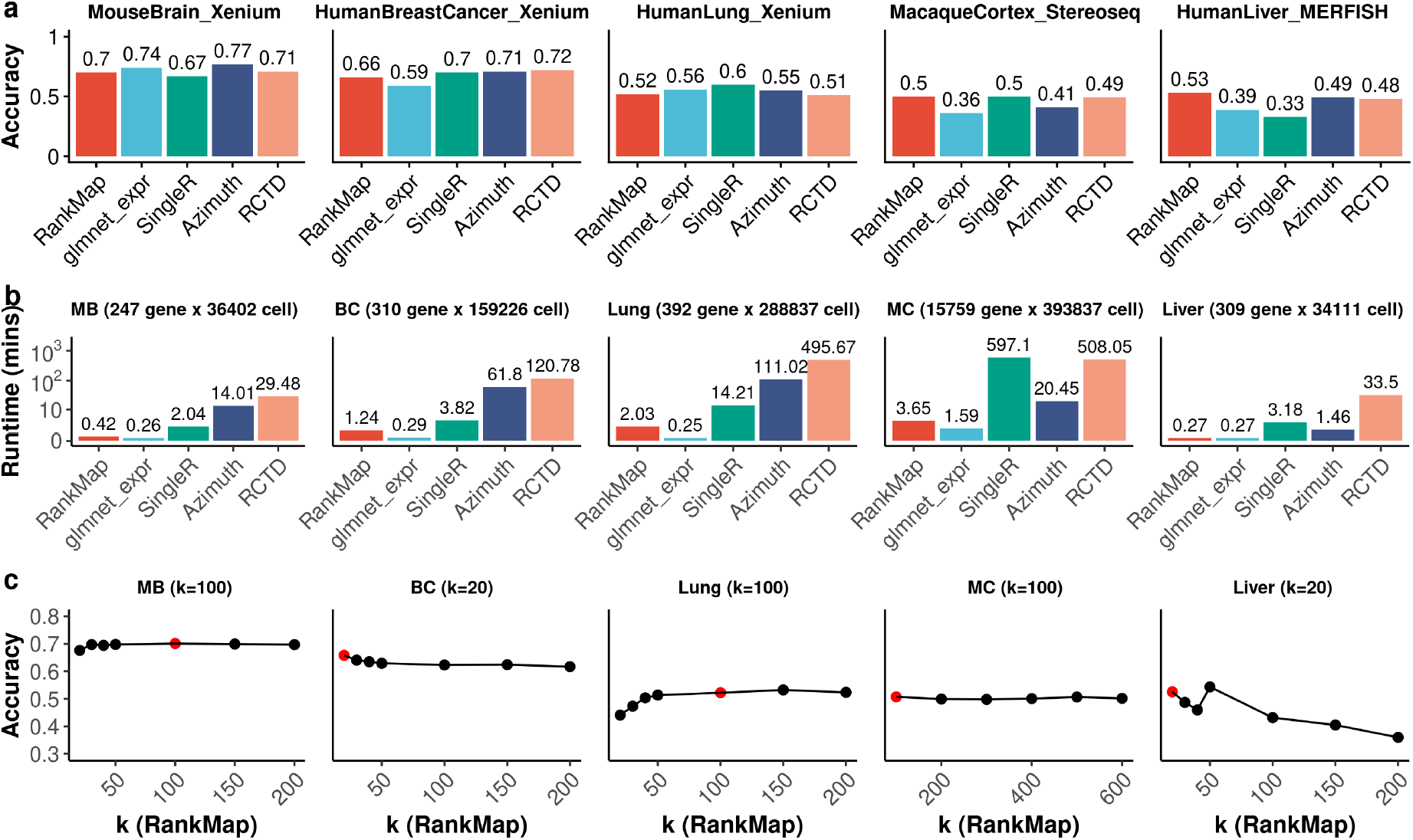
Benchmarking RankMap and other reference-based methods on spatial transcriptomics datasets. **a.** Accuracy of five reference-based cell type annotation methods (RankMap, glmnet_expr, SingleR, Azimuth, and RCTD) across five spatial transcriptomics datasets. Ground truth labels were derived from manual annotations (mouse brain, human breast cancer, human lung) or author-provided annotations (macaque cortex, human liver). Accuracy was calculated as the proportion of predicted labels matching the ground truth. **b**. Runtime (in minutes) of each method on the same spatial datasets. Gene and cell numbers are indicated for each dataset: mouse brain Xenium (247 genes, 36,402 cells), human breast cancer Xenium (310 genes, 159,226 cells), human lung Xenium (392 genes, 288,837 cells), macaque cortex Stereo-seq (15,759 genes, 393,837 cells), and human liver MERFISH (309 genes, 34,111 cells). RankMap automatically selects the top 500 variable genes when the number of common genes exceeds 500. **c**. Effect of the top-*k* parameter on RankMap accuracy. Line plots show classification accuracy across a range of *k* values for each spatial dataset. Red dots indicate the default *k* values used in this study (100 for mouse brain, human lung, and macaque cortex; 20 for human breast cancer and human liver). Optimal *k* values vary by dataset, reflecting differences in expression depth and cell type diversity.

In terms of runtime, *RankMap* was consistently faster than all other methods across the five spatial datasets (Figure 2b). On the mouse brain Xenium dataset, it completed predictions in 0.42 minutes, compared to 2.04 minutes for *SingleR*, 14.01 minutes for *Azimuth*, and 29.48 minutes for *RCTD*. On the breast cancer Xenium dataset, runtimes were 1.24, 3.82, 61.80, and 120.78 minutes, respectively. For the human lung dataset, runtimes were 2.03, 14.21, 111.02, and 495.67 minutes. For the macaque cortex dataset, runtimes were 3.65, 597.10, 20.45, and 508.05 minutes. For the liver MERFISH dataset, runtimes were 0.27, 3.18, 1.46, and 33.50 minutes. Across all datasets, *RankMap* was 3*×* to 244*×* faster than *Azimuth, SingleR*, and *RCTD*, with the largest speedups observed on large-scale datasets such as lung Xenium and Stereo-seq.

Together, these results demonstrate that *RankMap* provides competitive or superior accuracy while offering significantly improved computational efficiency across diverse spatial platforms and tissue types.

### 3.3 Evaluation of RankMap performance across different k values

The `k` parameter in *RankMap* determines how many of the top-expressed genes per cell are retained and ranked during feature transformation. This choice plays a crucial role in balancing signal retention and noise suppression. In our benchmarking analysis, we used `k = 100` for mouse brain, human lung, and macaque cortex datasets, and a smaller `k = 20` for human HER2+ breast cancer and human liver. The rationale for using smaller `k` values in the latter datasets is that closely related cell types, such as tumor cells, ductal carcinoma in situ (DCIS), and mature luminal (ML) cells in breast cancer, or the three hepatocyte subtypes in liver, often exhibit similar expression profiles that may be confounded by less informative genes.

To systematically evaluate the influence of `k` on performance, we tested a range of `k` values. For the Xenium and MERFISH datasets (each with a few hundred genes), we explored `k = 20, 30, 40, 50, 100, 150, 200`. For the Stereo-seq dataset (with whole-transcriptome coverage), we tested `k = 100, 200, 300, 400, 500, 600`. Prediction accuracy for each `k` value is shown in Figure 2c.

Across datasets, we found that accuracy generally improved as `k` increased up to a point, after which performance plateaued or declined. In the mouse brain and human lung datasets, accuracy increased with larger `k` values and stabilized around `k = 100-150`. For the macaque cortex dataset, which contains whole-transcriptome gene coverage, performance was stable across all tested `k` values, suggesting robustness to `k` selection in such contexts. In contrast, for the breast cancer and liver datasets, smaller `k` values (around 20–30) yielded the highest accuracy, while increasing `k` beyond 50 led to a gradual decrease. This decline is likely due to noise introduced by including additional genes that do not strongly distinguish among biologically similar cell types.

Overall, these results suggest that optimal `k` values may vary depending on the platform’s gene coverage and biological complexity. In general, moderate `k` values (e.g., 100) perform well for whole-transcriptome data, while smaller `k` values may be more suitable for panel-based data or samples with highly similar cell types.

### 3.4 Comparison of spatial organization of predicted cell types

To evaluate differences in spatial organization, we examined spatial maps of cell-type predictions generated by each method across the five spatial transcriptomics datasets (Fig. 3). These spatial visualizations, along with cell-type composition and method similarity analyses (Supplementary Fig. 1), provide insights into each method’s ability to recapitulate known anatomical and histological structures.

**Fig. 3.**
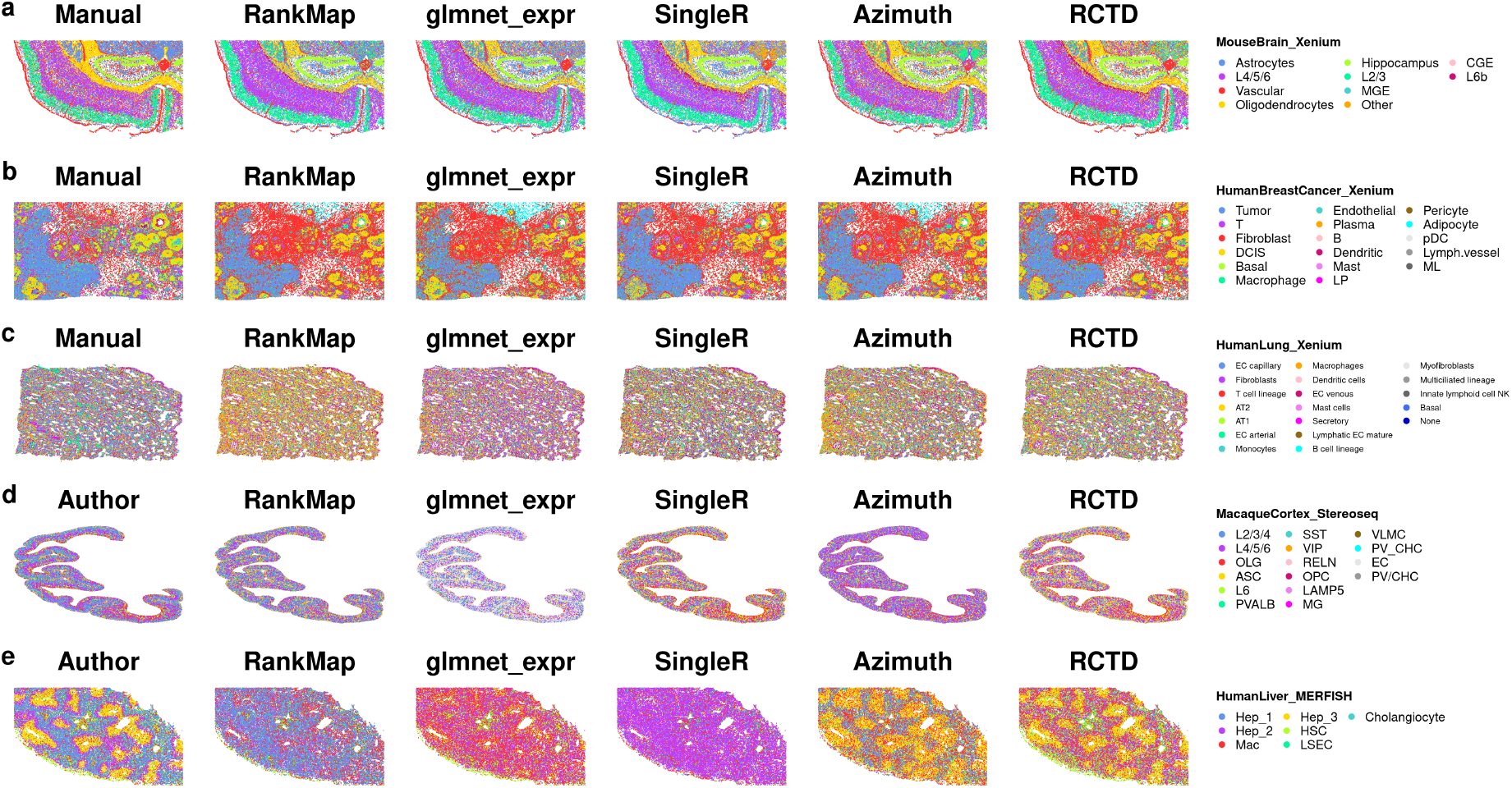
Spatial visualization of predicted cell types across datasets and annotation methods. **a.**Mouse brain (Xenium). **b**. Human breast cancer (Xenium). **c**. Human lung (Xenium). **d**. Macaque cortex (Stereo-seq). **e**. Human liver (MERFISH). Each panel displays spatial plots colored by predicted cell types from five reference-based annotation methods: RankMap, glmnet_expr, SingleR, Azimuth, and RCTD. Ground truth labels, derived from manual annotation (mouse brain, breast cancer, lung) or author-provided annotation (macaque cortex, liver), are shown for comparison. Visual differences highlight variation in spatial resolution, boundary preservation, and accuracy across methods and datasets.

In the mouse brain dataset, coronal sections reveal well-characterized anatomical regions including the meninges, cerebral cortex, hippocampal formation, and major white matter tracts [3, 15, 21, 22]. All methods, including manual annotation and the four reference-based tools, successfully captured the spatially coherent distribution of key cell types such as vascular cells in the meninges, layer-specific cortical neurons (L2/3, L4/5/6, L6b), hippocampal neurons, and oligodendrocytes in white matter (Fig. 3a). Cell-type composition across methods was highly concordant, and pairwise similarity metrics also indicated strong agreement (Supplementary Fig. 1a–b).

In the human HER2+ breast cancer dataset, tumor cells and DCIS cells are known to share similar gene expression profiles characteristic of the mature luminal (ML) lineage [1, 12, 16]. Spatial maps indicated that both cell types were correctly localized by all methods (Fig. 3b). Notably, an earlier benchmarking study reported frequent misclassification of tumor cells as DCIS by *Azimuth* when using the full reference dataset [12], whereas this issue was mitigated when using the 5k reference subset in the current study. Fibroblast predictions were consistent across methods but occurred at higher frequencies than in the manual annotations (Supplementary Fig. 1a). Overall, the pairwise similarity between methods remained high (Supplementary Fig. 1b).

For the human lung dataset, spatial predictions from *RankMap, SingleR, Azimuth*, and *RCTD* were broadly consistent with each other but showed differences compared to manual annotation (Fig. 3c). In particular, EC capillary cells, which were the most abundant cell type in the manual annotation, were underrepresented across all four reference-based methods (Supplementary Fig. 1). This suggests that distinguishing EC subtypes may be more challenging for automated label transfer approaches.

In the macaque cortex dataset, *RankMap* generated spatial distributions most similar to the author-provided annotation, whereas *glmnet_expr* produced the least similar map, predicting an overwhelming abundance of EC cells inconsistent with ground truth (Fig. 3d). *SingleR* and *RCTD* yielded similar results, while *Azimuth* showed modest deviations. Cell-type composition analysis reinforced the finding that *glmnet_expr* may not be suitable for this dataset due to its overprediction of EC cells (Supplementary Fig. 1).

In the liver MERFISH dataset, hepatocytes are expected to follow a zonated distribution across three liver lobule zones—periportal (Hep_1), midlobular (Hep_2), and pericentral (Hep_3)—based on their metabolic and transcriptomic profiles [19, 23–25]. The author-provided annotation accurately captured this zonation, with Hep_1 being the most abundant. However, the reference-based methods struggled to resolve the three hepatocyte subtypes (Fig. 3e). *RankMap* predominantly predicted Hep_1, *glm-net expr* misclassified most hepatocytes as macrophages, *SingleR* favored Hep_2, while *Azimuth* and *RCTD* primarily predicted Hep_3. Despite this variability, *RankMap* showed the highest agreement with the author-provided annotation, suggesting greater robustness in complex or partially overlapping cell-type scenarios.

Taken together, spatial visualization and comparative analysis reveal that *RankMap* produces spatially coherent and biologically plausible annotations across diverse datasets, often aligning more closely with expert annotations than other reference-based methods.

### 3.5 Benchmarking RankMap on single-cell datasets

To evaluate the performance of *RankMap* on single-cell data, we benchmarked it against two widely used reference-based annotation tools: *SingleR* and *Azimuth*. To minimize sample-specific bias and assess generalizability, we performed benchmarking across 12 human ER+ breast cancer samples and 8 healthy human lung samples. Accuracy and runtime metrics were computed for each individual sample.

Accuracy results are summarized in a bar plot (Fig. 4a). On the human ER+ breast cancer dataset, *RankMap* achieved accuracies ranging from 0.605 to 0.991, with a mean accuracy of 0.839. In contrast, *SingleR* showed a wider variance in performance, with accuracies ranging from 0.166 to 0.948 and a mean of 0.635. *Azimuth* showed intermediate performance, with accuracies ranging from 0.185 to 0.983 and a mean of 0.758. For the healthy lung samples, all three methods performed consistently well, with mean accuracies of 0.968 (*RankMap*), 0.969 (*SingleR*), and 0.968 (*Azimuth*).

**Fig. 4.**
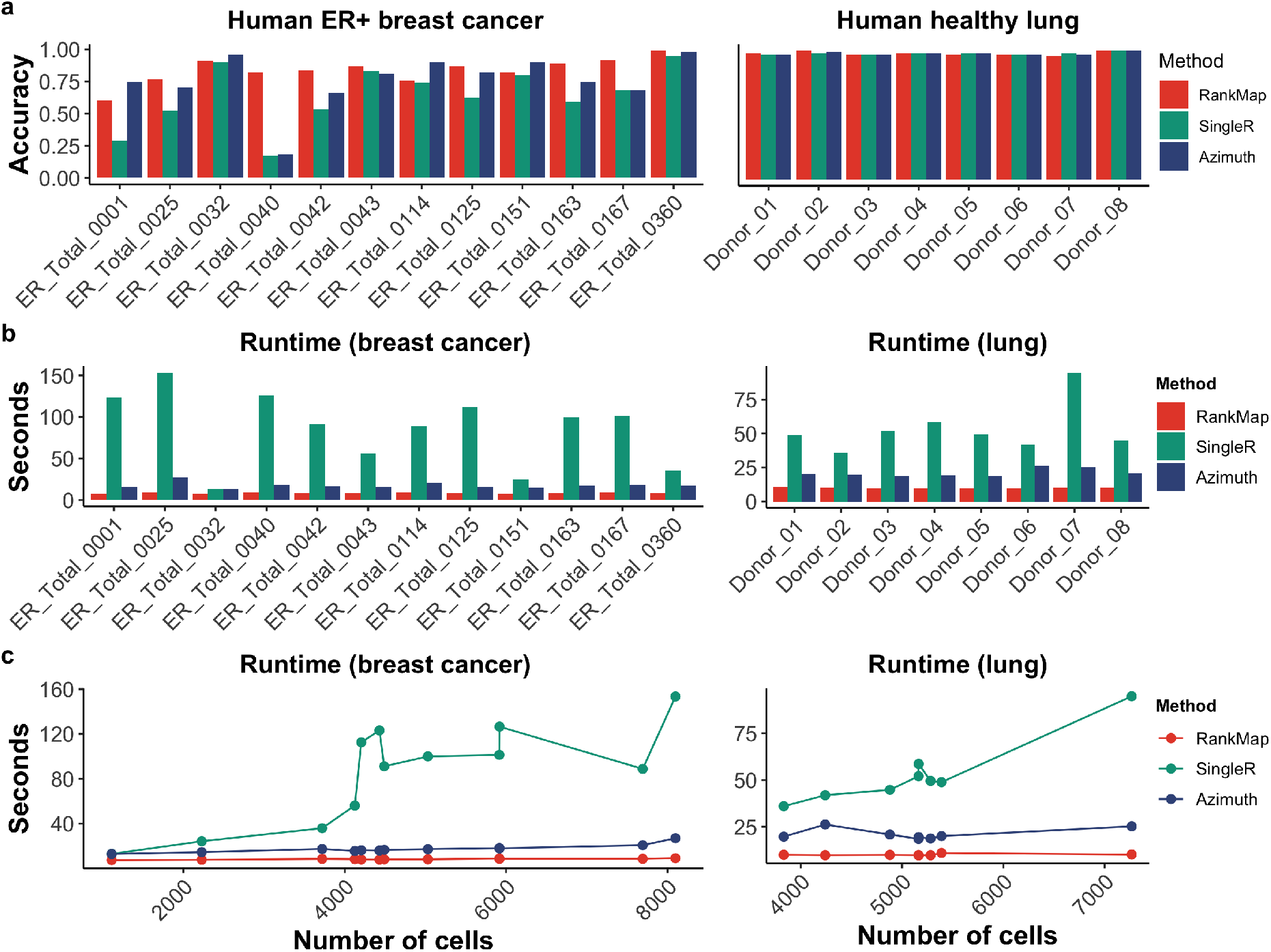
Benchmarking RankMap and other reference-based annotation methods on single-cell datasets. **a.**Accuracy of RankMap, SingleR, and Azimuth across 12 human ER+ breast cancer samples and 8 healthy human lung samples. Ground truth cell type labels were derived from manual annotations. One representative sample (ER_Total_0360 for breast cancer; Donor_02 for lung) was used as the reference to predict cell types in all other samples within each tissue type. **b**. Runtime in seconds for each method across the same single-cell datasets. **c**. Runtime for each method as a function of the number of query cells, illustrating scalability.

Runtime comparisons are shown in Fig. 4b. On the breast cancer samples, *RankMap* was the fastest method, with runtimes ranging from 7.51 to 9.26 seconds (mean: 8.39s). *Azimuth* followed with runtimes ranging from 12.99 to 27.07 seconds (mean: 17.62s), while *SingleR* was the slowest, ranging from 13.09 to 153.42 seconds (mean: 85.57s). A similar trend was observed for the lung dataset: *RankMap* had runtimes between 9.74 and 10.97 seconds (mean: 10.03s), compared to *Azimuth* (18.54–26.22s; mean: 21.10s) and *SingleR* (36.01–94.84s; mean: 53.34s). The runtime trend is further illustrated in Fig. 4c, which shows that *SingleR* runtime increases substantially with the number of cells in the query dataset, whereas *RankMap* and *Azimuth* are more scalable and less sensitive to query size. *RankMap* consistently outperformed both methods in speed across all samples.

Among the 12 breast cancer samples, two outliers (ER_Total_0040 and ER_Total_0001) exhibited particularly poor annotation performance with *SingleR*. To investigate these cases further, we examined UMAP visualizations of the predicted labels from each method alongside the reference sample ER_Total_0360 (Fig. 5). When ER_Total_0360 was used as both query and reference, all methods accurately annotated cell populations. However, for ER_Total_0040, *RankMap* correctly identified tumor cells, while *SingleR* and *Azimuth* misclassified many of them as ML cells. In ER Total 0001, both *RankMap* and *Azimuth* detected the tumor population, although some tumor cells were still labeled as ML. In contrast, *SingleR* failed to distinguish tumor from ML cells, misclassifying most tumor cells. UMAP plots for the remaining samples also showed superior separation of tumor and ML cell types by *RankMap* (Supplementary Fig. 2). In the lung dataset, all three methods performed well and correctly identified the expected cell types (Supplementary Fig. 3).

**Fig. 5.**
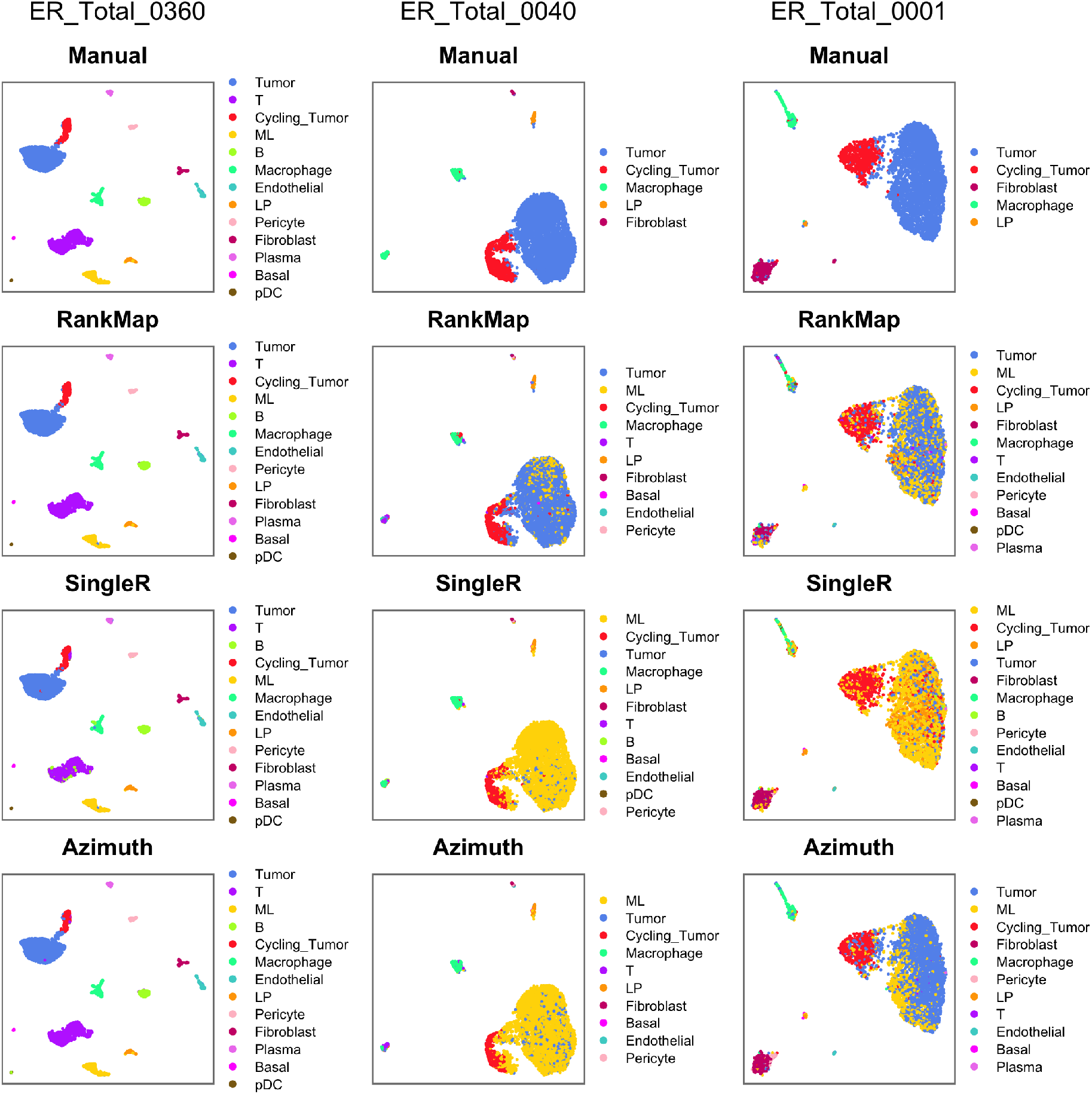
UMAP plots of selected human ER+ breast cancer single-cell samples colored by predicted cell types. Samples ER_Total_0360, ER_Total_0040, and ER_Total_0001 were chosen to illustrate the performance of RankMap, SingleR, and Azimuth. Sample ER_Total_0360 was used as the reference for predicting cell types in both itself and the other two samples. Manual annotations were used as the ground truth for comparison.

Overall, *RankMap* demonstrated the highest annotation accuracy and fastest runtime on the ER+ breast cancer dataset, while performing comparably to existing methods on the healthy lung dataset. These results highlight the robustness and efficiency of *RankMap*, particularly in complex or heterogeneous single-cell datasets.

## 4 Discussion

In this study, we introduced *RankMap*, a fast and flexible reference-based annotation method designed to support both single-cell and spatial transcriptomics data. By transforming gene expression profiles into ranked matrices and leveraging multinomial regression via *glmnet, RankMap* provides an efficient and robust approach to cell type annotation that is particularly well suited to modern spatial platforms.

Our benchmarking results demonstrate that *RankMap* achieves competitive or superior annotation accuracy across a diverse panel of datasets, including five spatial transcriptomics datasets (Xenium, Stereo-seq, MERFISH) and two single-cell datasets (ER+ breast cancer and healthy lung). Notably, *RankMap* consistently outperformed existing methods in runtime, completing predictions in seconds to minutes even on large spatial datasets with hundreds of thousands of cells. This scalability is a key advantage over tools such as *SingleR, Azimuth*, and *RCTD*, whose computational cost often scales poorly with dataset size.

The rank-based transformation confers several practical benefits. First, it mitigates batch effects and cross-platform variability, which are common when integrating spatial and scRNA-seq data. Second, it enables annotation using partial transcriptomes, making it well suited for spatial technologies with limited gene panels such as Xenium and MERFISH. Third, *RankMap* operates on commonly used data structures such as `Seurat`, `SingleCellExperiment`, and `SpatialExperiment`, simplifying integration into existing pipelines.

Despite these advantages, *RankMap* has several limitations. As a supervised reference-based method, its performance depends on the quality and representativeness of the reference dataset. In cases where the query data contain novel or rare cell states not present in the reference, the model may misclassify or overgeneralize. Recent generative-model-based tools, such as *CellDiffusion* [26], which annotate single-cell and spatial transcriptomics data using bulk references, have shown strong performance in annotating immune cell types and offer a promising complementary strategy to reference-based methods like *RankMap*. Additionally, although rank transformation improves robustness, it may obscure subtle transcriptional differences between closely related cell types, as observed in some breast cancer and hepatocyte subpopulations. The choice of the `k` parameter also influences performance, though we show that *RankMap* is generally stable across a range of `k` values.

Future work will focus on extending *RankMap* to incorporate additional contextual information. For example, integrating spatial coordinates or local neighborhood composition could refine predictions in heterogeneous tissues. Another promising direction is to allow multi-label or probabilistic annotations, particularly useful for transitional or mixed cell states. Finally, as spatial multi-omics data become more widely available, *RankMap*’s rank-based strategy may be adapted to incorporate non-transcriptomic modalities.

In conclusion, *RankMap* provides a robust, efficient, and easy-to-use solution for reference-based cell type annotation across both single-cell and spatial datasets. Its speed, generalizability, and compatibility with partial gene panels make it particularly valuable for large-scale spatial biology applications, where scalable and reproducible annotations are essential.

## Acknowledgements

We thank ChatGPT-4o (OpenAI) for the assistance with code development and documentation, as well as language editing. We also thank Yidi Sun and Juan Meng for sharing their published macaque cortex Stereo-seq spatial transcriptomics data (slide T33).

## Author’s contribution

J.C. conceived the RankMap package, collected data, conducted data analyses, interpreted results, generated figures and wrote the manuscript. S.L. collected data, discussed the analysis and interpreted results. S.K. collected data. C.H.A. and S.C.C. interpreted results. P.K.H.C and N.L. discussed and supervised this study. All authors read and approved the final manuscript.

## Availability of data and materials

In this study, publicly available datasets were used as detailed in Methods. Mouse brain tiny Xenium dataset: https://www.10xgenomics.comdatasets/fresh-frozen-mouse-brain-for-xenium-explorer-demo-1-standard. Mouse brain scRNA-seq dataset: GSE185862 [15]. Human breast cancer Xenium dataset and matched snRNA-seq dataset: https://www.10xgenomics.comproducts/xenium-in-situ/preview-dataset-human-breast[1]. Cheng et al annotation for the human breast cancer Xenium and snRNA-seq data: https://github.com/jinming-cheng/cheng_annotation_bmc_bioinfo [12] Human ER+ breast cancer scRNA-seq datasets: https://www.ncbi.nlm.nih.gov/geo/query/acc.cgi?acc=GSE161529[16]. Human healthy lung Xenium dataset: https://www.10xgenomics.com/datasets/xenium-human-lung-preview-data-1-standard. Human healthy lung scRNA-seq datasets (core): https://cellxgene.cziscience.com/collections/6f6d381a-7701-4781-935c-db10d30de293[17]. Macaque cortex Stereo-seq dataset and snRNA-seq dataset: https://www.braindatacenter.cn/datacenter/web/#/dataSet/details?id=1663381185152036865[18]. Human healthy liver MERFISH dataset and matched snRNA-seq dataset: https://datadryad.org/stash/dataset/doi:10.5061/dryad.37pvmcvsg[19].

## Code availability

The RankMap package is available at this GitHub repository: https://github.comjinming-cheng/RankMap. It is also available on Zenodo: https://zenodo.org/records18831050 [27]. Source code for reproducing the figures and analyses in this manuscript is publicly available at: https://github.com/jinming-cheng/Rcodes_for_RankMap_ms.

## Ethics approval and consent to participate

Not applicable.

## Consent for publication

Not applicable.

## 7 Supplementary Table and Figures

**Supplementary Table. 1.**
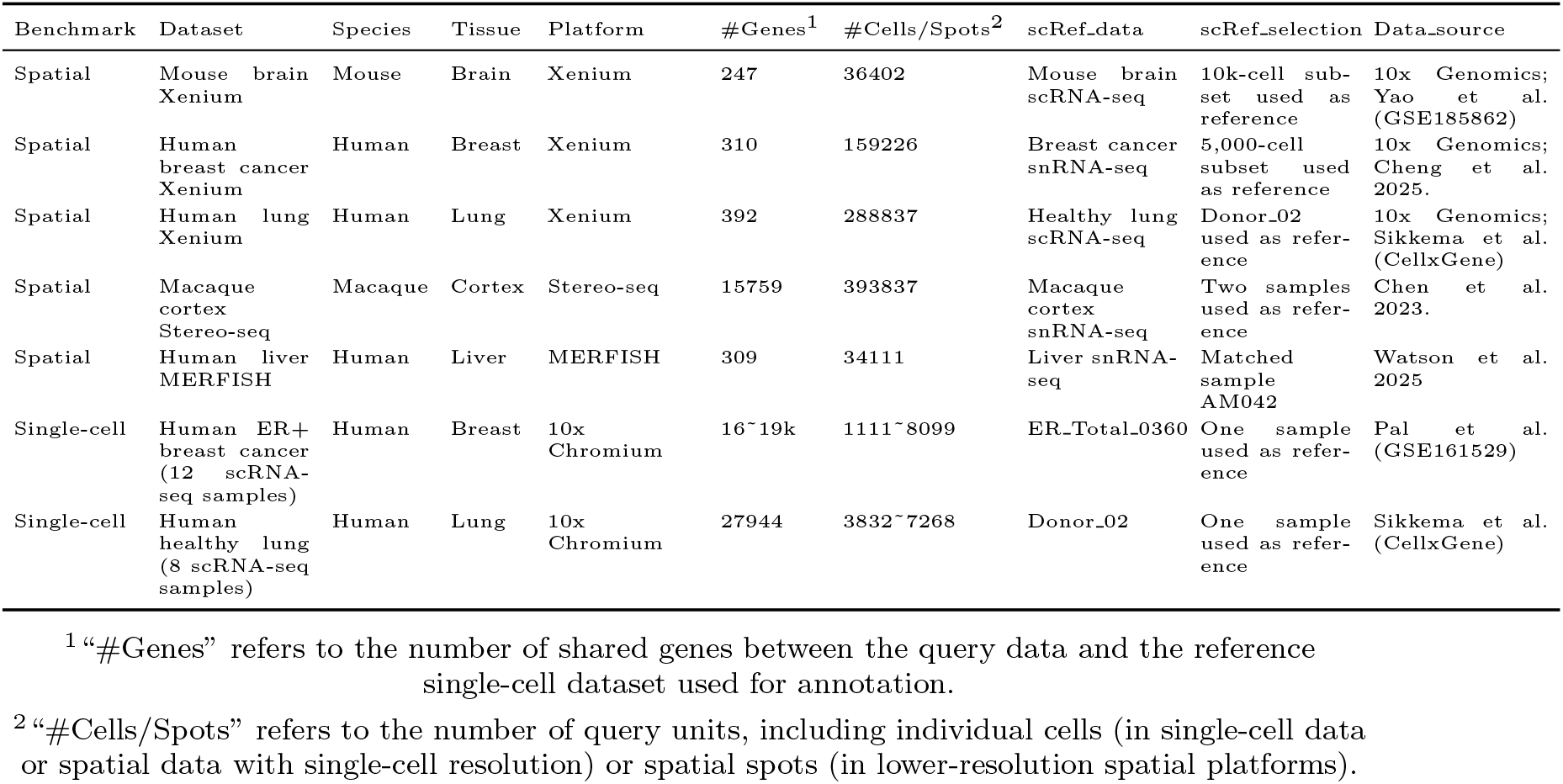
Summary of spatial and single-cell transcriptomics datasets used for benchmarking.

**Supplementary Fig. 1.**
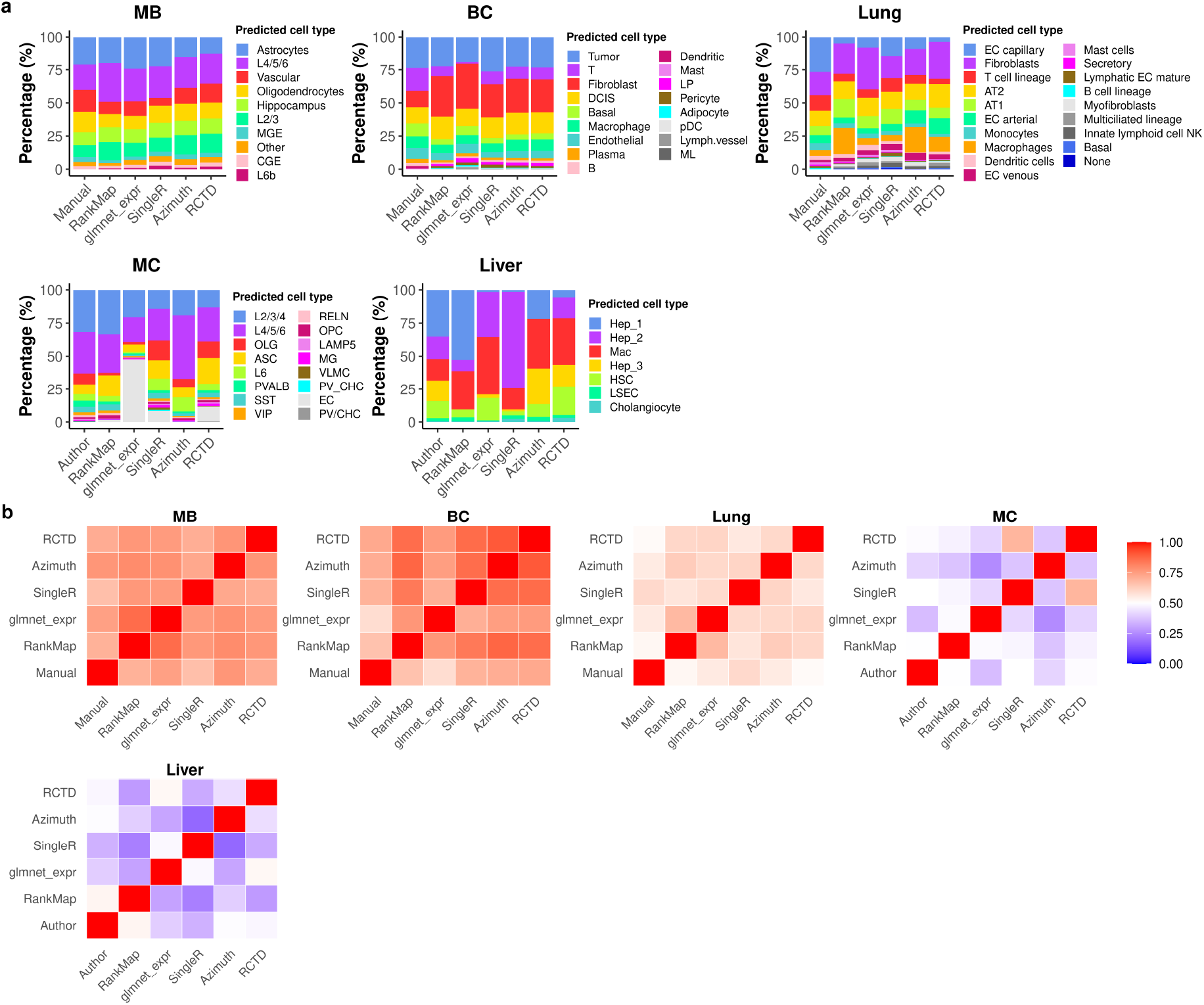
Comparison of cell type annotations across methods on spatial transcriptomics datasets. **a.** Bar plots showing the proportion of predicted cell types for each method across different spatial datasets. These plots highlight differences in cell type composition as assigned by each method. **b**. Heatmaps showing the pairwise agreement between methods for each spatial dataset, quantified as the percentage of shared predicted cell types.

**Supplementary Fig. 2.**
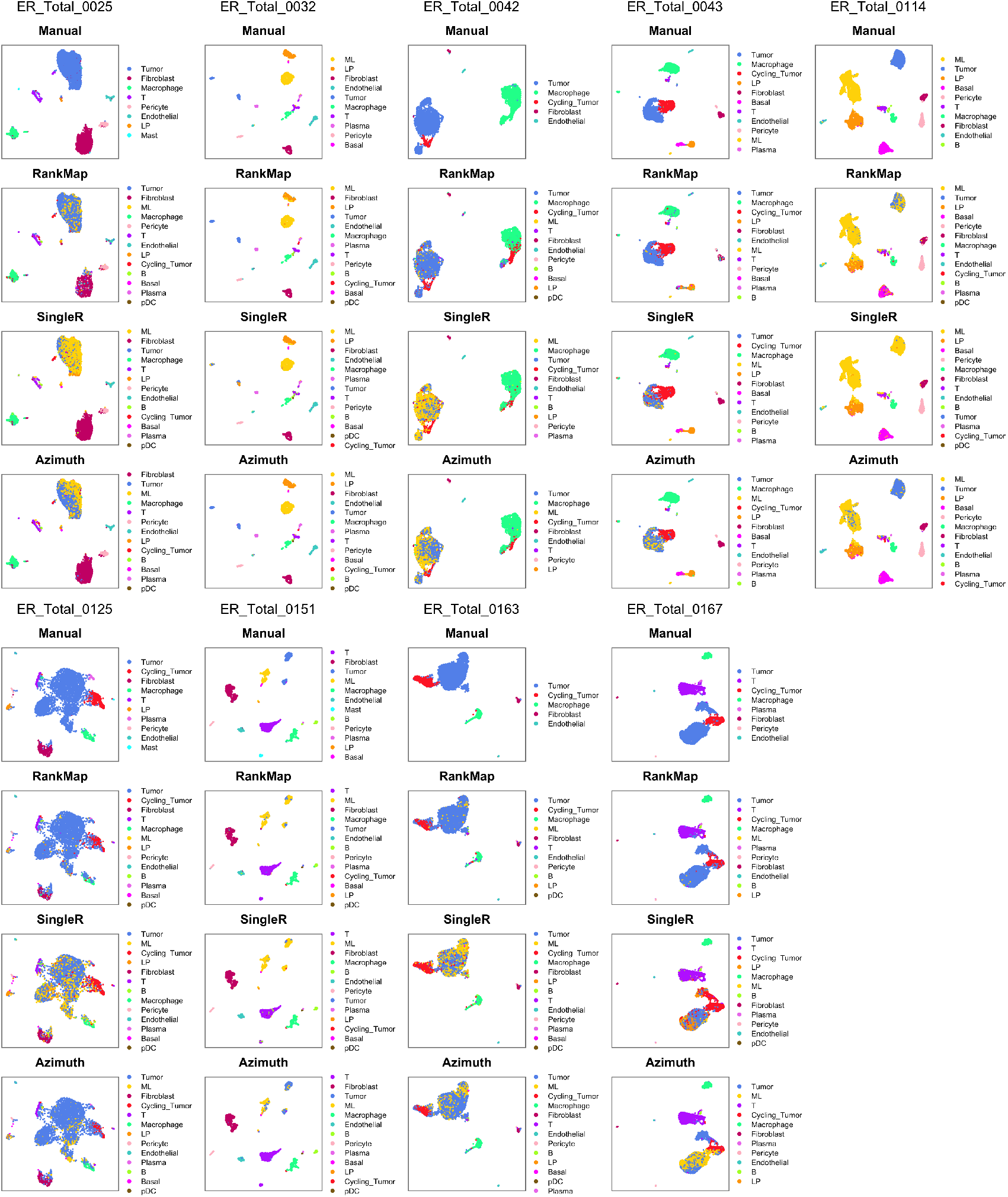
UMAP visualization of predicted cell types in additional human ER+ breast cancer single-cell samples. UMAP plots show cell type predictions for the remaining nine ER+ breast cancer single-cell samples. Predictions were made using RankMap, SingleR, and Azimuth, with sample ER Total 0360 serving as the reference. Manual annotations are provided as the ground truth for comparison. The plots illustrate method-specific differences in cell type resolution and consistency across samples.

**Supplementary Fig. 3.**
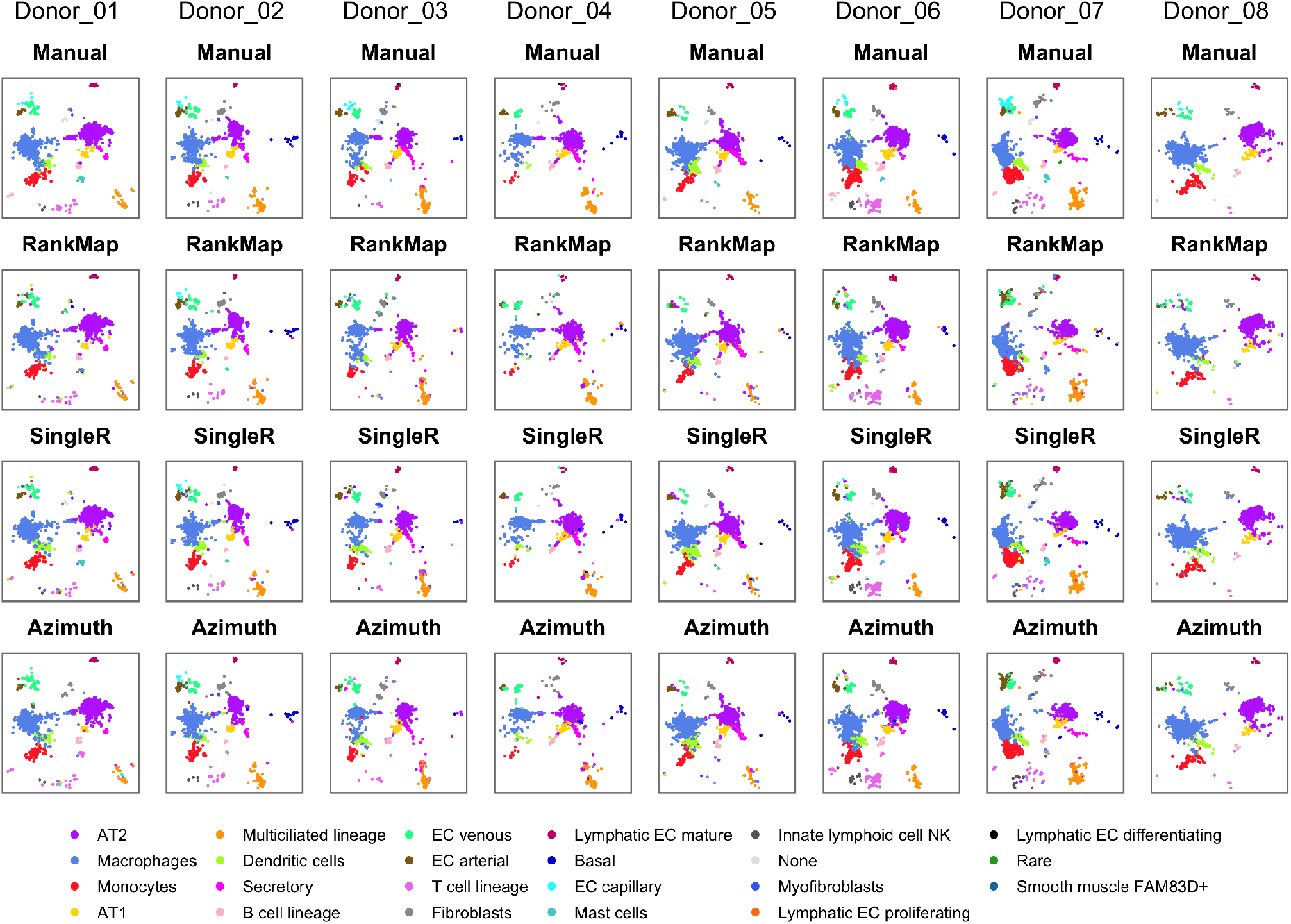
UMAP visualization of predicted cell types in human healthy lung single-cell datasets. Eight human lung samples are shown, with cell types predicted by reference-based annotation methods. Sample Donor 02 was used as the reference for predicting cell types in itself and the remaining samples. Each plot is colored by predicted cell types to highlight consistency and variability in annotation across samples.

